# CLUES2 Companion: Computational pipelines to estimate, visualize, and date selection on multi-locus sites

**DOI:** 10.1101/2025.06.10.658975

**Authors:** Alessandro Lisi, Michael C. Campbell

**Affiliations:** Department of Biological Sciences (Human and Evolutionary Biology Section), University of Southern California, 3616 Trousdale Parkway, Los Angeles, CA 90089

**Keywords:** selection coefficient, age of selection onset, allele trajectory, lactase persistence, natural selection

## Abstract

**Summary:** Statistical methods that quantify the selection coefficient associated with alleles provide critical insights into the evolutionary processes underlying organismal adaptation. Among these approaches, CLUES2 was recently developed to estimate selection coefficients for alleles using a statistical framework that captures the maximum amount of information present in genomic data, making it a state-of-the-art method for identifying variation under selection. However, before executing this approach, users first need to apply Relate, a genealogy-based approach, to their data to generate the input files for CLUES2. Moreover, completing this pre-processing step inherently assumes that users have sufficient expertise to successfully run the Relate software. Here, we present the CLUES2 Companion package, which contains user-friendly pipelines that seamlessly apply Relate to multiple sites within a target genomic region and then execute the CLUES2 software to estimate selection coefficients for these sites. CLUES2 Companion also has the capability to present the output of CLUES2 analyses in tabular and graphical formats. In addition, as a new feature, we adapted Relate and CLUES2 to estimate the age of onset of a selective sweep of derived variation, expanding the functionality of our package.

**Results:** To demonstrate the utility of our approach, we applied CLUES2 Companion to polymorphisms in the *MCM6* gene on Chromosome 2 (including the known variants associated with lactase persistence) in the European Finnish, Middle Eastern Bedouin, and East African Maasai populations from the 1000 Genomes Project, the Human Genome Diversity Project (HGDP), and the haplotype map (HapMap) Project Phase 3, respectively. Our analyses uncovered significant selection coefficient estimates at the persistence-associated T_−13910_ allele (rs4988235; s = 0.09986, CI: 0.08678 – 0.11294) in the Finnish, the G_-13915_ allele (rs41380347; s = 0.09981, CI: 0.06515 – 0.13448) in the Bedouin, and the C_-14010_ allele (rs145946881; s = 0.09981, CI: 0.08799 – 0.11163) in the Maasai, indicative of a classic selective sweep. Furthermore, we inferred the age of onset of selection at these alleles to be 9,100 years ago (CI: 6,552 – 10,612 years ago) in the Finnish, 7,700 years ago (CI: 1,864 – 8,064 years ago) in the Bedouin, and 4,900 years ago (CI: 3,864 – 5,936 years ago) in the Maasai, respectively, which coincide well with other estimates based on genetic and archaeological data. To further validate our dating method, we simulated several datasets containing SNPs with known ages of onset of selection, *s* estimates, and genomic positions using a selective sweep framework implemented in msprime and then applied CLUES2 Companion to the simulated datasets. Using this approach, CLUES2 Companion produced similar estimates of selection onset as the ones specified in the simulations, corroborating the dependability of our method. Overall, CLUES2 Companion is a versatile package that enables users to efficiently explore, interpret, and report evidence of selection in genomic datasets, complementing the CLUES2 software.

**Availability and Implementation:** CLUES2 Companion is free and open source on GitHub (https://github.com/alisi1989/CLUES2-Companion) and on DropBox (https://www.dropbox.com/scl/fo/m5y6aek0twd1jz9grg4p3/ALxMgIljUJRIZZNQXaGU-OE?rlkey=mbbh36ondftnqg0×07eao57eg&st=9jzsl5um&dl=0).

## Introduction

The selection coefficient, *s*, quantifies the fitness effects of mutations that arise in populations, providing insight into the impact of selection at the genetic level (Thurman and Barrett 2016; Xu, Futschik and Dutta 2022; Nielsen, Vaughn and Deng 2025; Zhao *et al*. 2025). To date, several statistical methods have been developed to infer *s* at genetic loci (Bollback, York and Nielsen 2008; Illingworth and Mustonen 2011; Illingworth *et al*. 2012; Malaspinas *et al*. 2012; Tamuri, dos Reis and Goldstein 2012; Mathieson and McVean 2013; Terhorst, Schlötterer and Song 2015; Topa *et al*. 2015; Schraiber, Evans and Slatkin 2016; Tataru *et al*. 2016; Iranmehr *et al*. 2017; Taus, Futschik and Schlötterer 2017; Zhao *et al*. 2017; Kirsch-Gerweck *et al*. 2023; Likhachev and Rouzine 2023; Sibly and Curnow 2023; Vaughn and Nielsen 2024). Among them, CLUES2 incorporates ancestral recombination graphs (ARGs)—which capture the history of genomic regions in a sample of sequences—along with importance sampling of ARGs to estimate *s* in different time periods (Vaughn and Nielsen 2024). Thus, CLUES2 is argued to be a state-of-the-art method that utilizes the full set of allele histories in a sample of sequences to reliably estimate *s* through time (Vaughn and Nielsen 2024).

However, before executing this approach, users first need to process their data with Relate (Speidel *et al*. 2019), a method for estimating genealogies, to generate the input files for CLUES2 (Vaughn and Nielsen 2024). More specifically, using Relate, individuals must: i) convert fully phased vcf file, containing single nuclear polymorphisms (SNPs), to *haps and *.sample files; ii) identify the derived allele at each SNP in the dataset (i.e., polarize alleles); iii) generate the *.anc and *.mut files, (containing polarized derived allele information) with a user-defined effective population size (*N*_e_); iv) create the poplabels file (*.poplabels); v) generate the coalescence file (*.coal); vi) update the *.anc and *.mut files using the *.coal file; vii) sample branch lengths using the *.anc, *.mut files, and *.coal files together with the start and end positions for each SNP of interest; and viii) generate a *.newick file for each SNP. Users then will apply the CLUES2 software, which consists of two Python scripts: i) RelatetoCLUES.py and ii) inference.py. RelatetoCLUES.py converts the *.newick and the polarized derived allele (*.txt) files from Relate to a *.times file. Inference.py will use the *.times, *.coal, and derived allele frequency files to estimate *s* for a given SNP. However, performing these intricate pre-processing steps assumes that users have sufficient expertise to successfully produce the required files for CLUES2 with the Relate software. Furthermore, because the steps described above are applied to SNPs one at a time, there is potential for users to introduce errors into their analyses if they are examining multiple SNPs.

To mitigate these challenges, we developed the CLUES2 Companion package that seamlessly creates the input files for multiple sites within a genomic region using Relate and then applies the CLUES2 software to estimate *s* for these loci (Vaughn and Nielsen 2024). CLUES2 Companion also has the capacity to display the numerical results of CLUES2 analyses as tabular and graphical outputs. To demonstrate the utility of our approach, we applied the CLUES2 Companion to the *MCM6* gene (36,818 bp in length) on Chromosome 2, which contains polymorphisms that contribute to lactase persistence (i.e. the ability to digest the lactose sugar in fresh milk) in the European Finnish, Middle Eastern Bedouin, and East African Maasai populations. Our analyses revealed that the derived alleles associated with lactase persistence— namely, the T_-13910_, G_-13915,_ and C_-14010_ alleles—exhibited significant *s* estimates indicative of a classic selective sweep at these loci. Furthermore, using the dating method implemented in CLUES2 Companion, we inferred the age of onset of selection for these functional alleles, which ranged from 4,900 to 9,100 years ago in agreement with prior genetic and archaeological studies. Overall, CLUES2 Companion is a versatile package that enables users to efficiently scan genomic regions for important functional variants interactively with the Relate and CLUES2 software (Vaughn and Nielsen 2024).

## 2. Materials and Methods

CLUES2 Companion consists of three distinct pipelines written mainly in Python: i) Phase 1 (which generates population-specific input files for Relate); ii) Phase 2 (which estimates *s* and confidence intervals along with *P*-values); and iii) Phase 3 (an optional step that infers the age of selection onset for SNPs with significant *s* values).

### 2.1 Processing phased vcf data to generate population-specific input files to run Relate (Phase 1)

Prior to executing Phase 1, users will need: i) fully phased variant call format (vcf) files with bi-allelic sites split by chromosome; ii) an ancestral allele FASTA file (located on our DropBox repository at https://www.dropbox.com/scl/fo/dgpho1vd736xo317osdem/AJUo5D4rq41LT2-YR5JGHeU?rlkey=76co7uit9f0dkkhxc7yqvb0kl&st=6y1ii6uj&dl=0 or at https://ftp.ensembl.org/pub/release112/fasta/ancestral_alleles/); iii) a pilot mask file (where alleles are coded as either high quality or low quality) found at https://www.dropbox.com/scl/fo/3v1zh09cezqob9vcazcpj/AAuPvFYt6SedvTumMV140pk?rlkey=5orcm01mbhczrvdtvt3m2p24h&st=yk6qtyoy&dl=0; iv) generic genetic maps formatted by Relate (https://www.dropbox.com/scl/fo/m5ftor1edz9rrykyfg7te/AHGuA84I6cDQgx5bhe6aiLo?rlkey=4k54o1bo79v11tqjywl8a9xeb&st=jtfrtipl&dl=0); and v) *.poplabels file that users must generate manually based on information in the Relate manual (https://myersgroup.github.io/relate/input_data.html); an example of the *.poplabels file can be found at https://github.com/alisi1989/CLUES2-Companion/tree/main/example).

With the above files and other information provided by users (e.g. chromosome number, start and end positions of a target region to analyze, etc.), Phase 1 will convert the *.vcf file to *.haps and *.sample files (which contain phased data and a list of sample names, respectively). Our Phase 1 pipeline will then infer the ancestral or derived state of alleles at each biallelic site using either the Homo_sapiens_hg38_reference FASTA file (Cunningham *et al*. 2022; Lisi and Campbell 2024) or the ancestral allele file provided by Relate (Speidel *et al*. 2019) as reference. After this polarization step, our pipeline will calculate derived allele counts and frequencies per site. Notably, the accurate polarization of alleles is critical since *s* is calculated using derived alleles. If this procedure is not performed correctly, users could inadvertently create new data. Phase 1 will also execute Relate to generate the coalescence (*.coal), tree information including branch lengths (*.anc), and SNP-specific information (*.mut) files.

In addition, our pipeline will automatically create a Phase 1 folder using the abspath function and save the following files to this folder: i) the count and frequency of the polarized derived alleles at SNPs within the specified target region (*.txt file); ii) the coded derived allele file for each SNP in the specified region (one coded allele per line; each line represents a sample); iii) SNP identifiers and associated genomic coordinates (*.txt file); iv) the *.anc, *.mut, and *.coal files; and v) the *.log file.

### 2.2 Creating the *newick files for each SNP using Relate and then applying CLUES2 to calculate *s* and the associated confidence intervals (Phase 2)

Like in Phase 1, Phase 2 will auto-detect the location of input files for this step. To activate this capability, users only need to enter the population prefix (e.g., Finnish) used in Phase 1 when prompted. Then, CLUES2 Companion will automatically retrieve the files from the Phase 1 folder and use them as input for Phase 2. The Phase 2 pipeline will: i) apply Relate to sample branch lengths using the *.coal file generated in Phase 1; and ii) produce a *.newick file for each SNP in a target region specified by users. Notably, our pipeline will simultaneously produce a *.newick file for each SNP within a region on a given chromosome. In comparison, if users applied the Relate software directly to their data, without CLUES2 Companion, the sampling of branch lengths would be applied separately to each SNP, resulting in a *.newick file for that SNP. That is to say, if users were interested in analyzing 100 SNPs, for example, they would need to apply the Relate software 100 times to generate a *.newick file for each SNP. Therefore, a major advantage of our Phase 2 pipeline is its ability to seamlessly process multiple SNPs within a genomic region with minimal error.

The CLUES2 software consists of two Python scripts: i) RelatetoCLUES.py and ii) inference.py. CLUES2 Companion will invoke CLUES2, first running the RelatetoCLUES.py script to create a *.times file for each SNP, which will serve as input for inference.py. Next, to execute inference.py (with the *.times file as input), users will be prompted to provide arguments for the following: i) df (the number of points used to bin the allele frequencies); ii) tCutoff (which is maximum time in generations to be considered for analysis); iii) importance sampling of branch lengths (number of times branch lengths are sampled); iv) AncientSamps and AncientHaps (if using ancient DNA data); v) noAlleleTraj (the inferred allele frequency trajectory); vi) ci (a list of confidence intervals, e.g. 0.95); and vii) --TimeBins (a list of epoch breakpoints).

Using this information, inference.py will then estimate *s* for each SNP, confidence intervals, *P*-values, and bin generations.

Additionally, Phase 2 will create a tabular output file in *.tsv format containing the rs identifiers of SNPs within a target region, their genomic positions, the derived allele frequency at each SNP, logLR, negative log10(p), epoch start, epoch end, *s* estimate (MLE1). and confidence intervals (**Fig. 1; Tables S1-S2**). Phase 2 will also generate a selection coefficient intensity plot of SNPs within a target region along with the rs identifiers for SNPs that exhibit significant s values *(P* < 0.05). Furthermore, this plot will show color-coded intensity based on the *P*–values, allowing users to immediately recognize the degree to which *s* estimates are significant in addition to the asterisks (**Fig. 1; Fig. S1**). Finally, Phase 2 will automatically create a Phase 2 folder to which all output files, including the *.log file, will be saved. These output files together with the output files from Phase 1 will serve as input files for Phase 3 (**Fig. 1; Fig. S1**).

**Figure 1.**
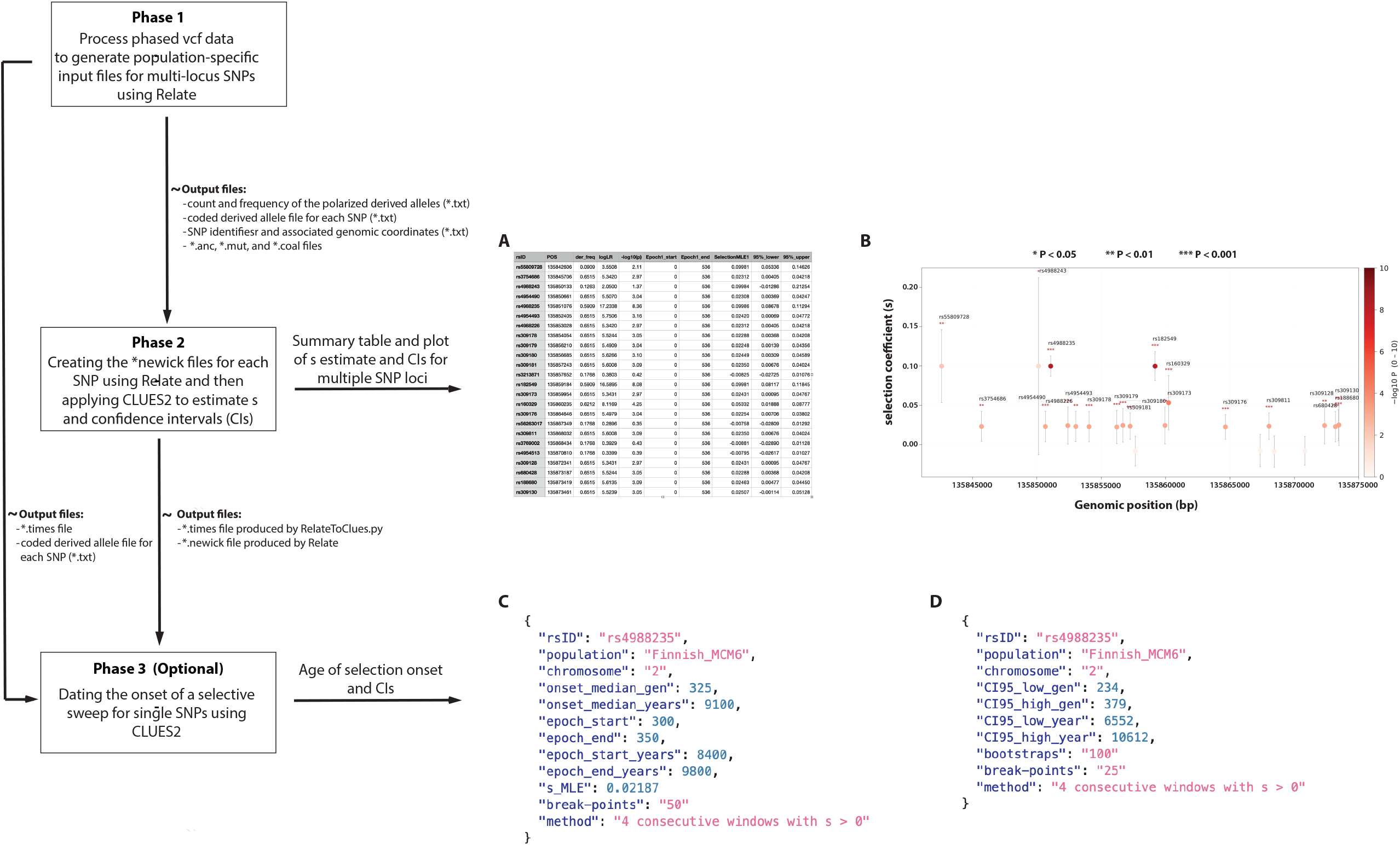
Workflow of the CLUES2 Companion package. CLUES2 Companion consists of three distinct pipelines: i) Phase 1, ii) Phase 2, and iii) Phase 3 that run in a command-line Terminal. The input for Phase 1 consists of fully phased vcf files split by chromosome containing bi-allelic sites. Using the output files from Phase 1, Phase 2 creates *.newick files (with tree information) for SNPs using Relate and then applies CLUES2 to calculate *s* and the associated confidence intervals. The Phase 2 pipeline can also generate the output of CLUES2 analyses in tabular and graphical formats. For example, **Panel A** shows the tabular output in *.tsv format containing the rs identifiers for SNPs in the *MCM6* gene (including the rs4988235 SNP associated with lactase persistence), their genomic positions, the derived allele frequency at each SNP, logLR, -log10(p), epoch start, epoch end, *s* estimate (MLE1) and confidence intervals in the Finnish population. **Panel B** displays a plot of *s* estimates for SNPs (above a frequency of 5%) in the *MCM6* gene in the Finnish together with the rs identifiers for SNPs that exhibit significant s values *(P* < 0.05). In the plot, each dot represents an *s* estimate, and the vertical bars indicate the confidence intervals around the estimate for each SNP (exact numbers can be found in the tabular *.tsv file). Furthermore, the color of each dot signifies the level of significance for each s estimate based on *P*–values in addition to the number of asterisks. Phase 3 is an optional step that infers the age of onset of a selective sweep for SNPs with significant *s* estimates using the input files from Phase 1 and Phase 2. For example, **Panel C** shows the age of onset of selection on the derived allele at the T_-13910_ allele (rs4988235) in generations and years (one generation=28 years) in the Finnish along with the median value of the oldest earliest window of time in which selection occurred, the window range, and the associated s estimates. **Panel D** depicts the 95% confidence intervals around the estimated age of selection on the T_-13910_ allele (rs4988235) in the Finnish and the method used to determine the lower and upper bounds of the confidence interval. The results of the Phase 2 and Phase 3 analyses for the Bedouin and Maasai can be found in the online Supplementary data.

### 2.3 Estimating the age of onset of a selective sweep for SNPs with significant *s* estimates (optional Phase 3)

We formulated an approach to infer the onset of selection on derived variation using the computational framework in Relate and CLUES2 (Speidel *et al*. 2019; Vaughn and Nielsen 2024). To begin, users will be prompted to provide the: i) chromosome number to analyze; ii) population prefix (the same as the prefix in Phase 2); iii) rs identifier (for the SNP to date); iv) df (the number of breakpoints used to bin the derived allele frequency; df=50 is the default); and v) tCutoff (the maximum time in generations to be considered in the analysis). Phase 3 will calculate the onset of selection for a given SNP using a series of temporal breakpoints (or windows) from 0 to 2,000 generations (gen.) ago. More explicitly, our pipeline will segment time into non-overlapping windows of size 50 gen. (e.g., 0–50 gen., 50–100 gen., 100–150 gen.) from 0 to 2,000 gen. ago. The default window size of 50 gen. was determined to be a reasonable balance between accurately calculating *s* and computational efficiency based on empirical testing. However, users have the option to choose their own window size.

Next, the Phase 3 pipeline will estimate *s* and associated *P*–value for each of the non-overlapping windows (50 gen.) from 0 to 2,000 gen. ago. Then, the age of onset of selection on derived alleles will be determined using a trend-based approach. Specifically, if four consecutive windows with *s* > 0 and *P* < 0.05 (or -log_10_(*p*) > 1.30103) are observed between 0 and 2,000 gen. ago in the past, we concluded that the onset of selection occurred in the oldest of the four consecutive windows (specifically, the median value of the oldest window), representing the earliest time (*t*_*o*_) of selection onset. The choice of four consecutive bins was determined based on empirical testing of our approach. If four consecutive windows exhibiting *s* > 0 and *P* < 0.05 between 0 and 2,000 gen. ago are not found, our pipeline then searches for three consecutive windows with these criteria; similarly, the onset of selection will be inferred to have occurred in the oldest of the three consecutive windows. If three consecutive windows are not present, our pipeline will search for two consecutive windows in which *s* > 0 and *P* < 0.05 (the onset of selection will be determined using the method described above). If these two consecutive windows do not exist, Phase 3 will print the following warning to the Terminal: “No two consecutive windows found. The window with s > 0 and the highest -log_10_(*p*) value of all scanned windows will be taken as the age of selection onset”. The resulting output files from these analyses will be automatically saved to the Phase 3 folder.

To calculate confidence intervals, users will be prompted to provide a start and end time around the estimated age of onset at a given SNP. For example, if the onset of selection were inferred to be 300 gen. ago, users could choose a lower bound of 0 and an upper bound of 2,000 generations ago, as a starting point, to calculate the confidence intervals. Alternatively, to decrease the computational time, users could choose a smaller range, for example a lower bound of 0 and upper bound of 500 gen. ago. Regardless of the choice, our method will calculate *s* for each segmented non-overlapping window within the specified range (25 gen. is the default window size) and scan for consecutive windows with *s* > 0 and *P* < 0.05 to infer time of selection onset, *t*_n_, using the same trend-based procedure described above. This process of running Relate to generate genealogies (Phase 1) and CLUES2 to calculate *s* and *P*–values in segmented windows (Phases 2 and 3) for a given SNP will be repeated *n* times (*n* the number of replicates defined by the user) with a random seed, generating an empirical distribution of *t*_n_ (e.g. *t*_1_…*t*_1000_). The Phase 3 pipeline then will calculate the 2.5^th^ and 97.5^th^ percentiles of this distribution, representing the lower and upper bounds of the confidence interval, respectively.

## 3 Results and Discussion

We developed user-friendly pipelines to: i) run the Relate software to create the input files for CLUES2; ii) estimate s and other statistics using CLUE2; iii) display the numerical output of this analysis in tabular and graphical formats; and iv) infer the age of selection onset of alleles with significant *s* estimates. To demonstrate the efficacity of our approaches, we applied CLUES2 Companion to genetic variation in the *MCM6* gene on Chromosome 2 in the European Finnish, Middle Eastern Bedouin, and East African Maasai populations from the 1000 Genomes Project, the Human Genome Diversity Project (HGDP), and the haplotype map (HapMap) Project Phase 3, respectively (International HapMap Consortium 2003; The 1000 Genomes Project Consortium 2012; Bergström et al. 2020). More explicitly, we estimated *s* for allelic variation in *MCM6*, including the functional T_−13910_, G_-13915_, and C_-14010_ alleles that play a role in lactase persistence and are known to be targets of selection (Enattah et al. 2002; Ingram et al. 2007, 2009; Tishkoff et al. 2007; Ranciaro et al. 2014; Hassan et al. 2016; AnguitaRuiz, Aguilera and Gil 2020; Campbell and Ranciaro 2021; Lisi and Campbell 2024). Our analyses uncovered significant *s* estimates (*P* < 0.05) at the derived T_−13910_ allele in the Finnish (rs4988235; s=0.09986, CI: 0.08678– 0.11294), the derived G_-13915_ allele in the Bedouin (rs41380347; s=0.09981, CI: 0.06515–0.13448), and the derived C_-14010_ allele in the Maasai (rs145946881; s=0.09981, CI: 0.08799–0.11163) populations, indicative of a classic selective sweep (**Fig. 1; Table S1; Figs. S1-S7**). Moreover, these estimates are comparable to those inferred in similar populations with a history of production and consumption (Tishkoff *et al*. 2007; Enattah *et al*. 2008; Ségurel and Bon 2017; Burger *et al*. 2020; Kirsch-Gerweck *et al*. 2023; Vaughn and Nielsen 2024). For example, Ennatah and colleagues (2008) reported that *s* estimates associated with the derived G_−13915_ allele varied between 0.04 and 0.14 in Middle Eastern and North African populations (Enattah *et al*. 2008; Ségurel and Bon 2017). Notably, our analyses also showed that the other alleles in *MCM6* with significant s values were in linkage disequilibrium (LD) with these known functional sites (**Tables S3-S5**).

In addition, we estimated the age of onset of selection for the T_−13910_ allele to be 9,100 years ago (CI: 6,552 – 10,612 years ago) in the Finnish, 7,700 years ago (CI: 1,864 – 8,064 years ago) for the G_–13915_ allele in the Bedouin, and 4,900 years ago (CI: 3,864 – 5,936 years ago) for the C_-14010_ allele in the Maasai. Prior studies also have reported similar time estimates for selection onset in milk-dependent populations (Loftus *et al*. 1994; Bersaglieri *et al*. 2004; Ségurel and Bon 2017; Roffet-Salque *et al*. 2018; Makarewicz 2020; Campbell and Ranciaro 2021; Alkaraki *et al*. 2024). For example, Bersaglieri et al. (2004) indicated that haplotypes carrying the T_-13910_ allele began to rise rapidly in frequency in populations of European descent between 2,188 and 20,650 years ago, coinciding with the estimated origin of dairy farming in northern Europe ∼9,000 years ago (Bersaglieri *et al*. 2004). To further validate our dating approach, we simulated several datasets with known parameters for target SNPs (such as age of selection onset, *s* estimate, genomic position) using the SweepGenicSelection and sim_ancestry functions in msprime (Baumdicker *et al*. 2022). We then applied our Phase 3 pipeline to these simulated data to see if our dating method would reproduce the ages of onset for the target SNPs in the simulated data. Using this procedure, our dating method produced similar estimates of selection onset as the ones assigned to alleles in the simulations (**Table S6**), corroborating the reliability of our approach.

In summary, CLUES2 Companion has the capacity to efficiently: i) estimate *s* and other statistics; iii) present the numerical output; and iv) infer the age of selection onset. While these strengths make CLUES2 Companion a powerful tool for investigating, interpreting, and reporting evidence of selection, it is also important to note the limitations of our methods. In particular, calculating the confidence intervals around the age of selection onset (*t*_*0*_) is computationally intensive because our pipeline iteratively searches for this time of origin in small, segmented windows across a large time scale multiple times. For example, with a window size of 25 gen. ago over a time scale ranging from 0 to 500 gen. ago, a df of 600, importance sampling of 200, and 100 bootstrap replicates, the compute time to calculate the confidence interval around an age estimate of 300 gen. ago for the T_-13910_ allele in the Finnish was between 16 and 24 hours on a Desktop computer with 24-core (2.4 GHz) node and 32 GB RAM. Furthermore, this runtime will increase if users choose to use a window size smaller than 25 gen. and a larger number of replicates (e.g., 1,000). While our approaches yield reliable results, some users may consider this compute time to be long. To mitigate this issue, we recommend that they consider applying a parallel multithreading approach on multiple cores (or engage multiple threads on a single core) to reduce the computing wait times. However, if this option is not available, users should decide for themselves how computationally intensive they wish to make their analyses (until we devise a way to reduce the runtime in a future iteration of our package).

Finally, although significant *s* estimates at loci within a genomic region are strongly indicative of selective sweeps, it is possible that not all of these loci are or have been the direct targets of selection. Instead, a subset of these alleles could exhibit significant s estimates because they are in LD with the actual allele(s) under selection (**Tables S3-S5**). While this issue is not directly related to CLUES2 Companion and CLUES2, it is worthwhile mentioning that users should corroborate signals of selection with additional lines of evidence like with any analysis. Nonetheless, if prudently applied, CLUES2 Companion is a highly beneficial tool that can assist with identifying and visualizing mutations that have contributed to organismal adaptation and evolution.

## Acknowledgements

We thank our novice and experienced users for testing the reproducibility of our computational pipelines. We also thank Professor Rasmus Nielsen and Dr. Andrew Vaughn for their feedback on our computational pipelines and for their critical review of the manuscript.

## Supplementary data

Supplementary data are available at *Bioinformatics* online.

## Data availability

The data underlying this article are available in the 1000 Genomes Consortium Project at https://www.internationalgenome.org/data-portal/population/FIN; in the HGDP at https://ngs.sanger.ac.uk/production/hgdp/hgdp_wgs.20190516/statphase/; and in the HapMap Project Phase 3 at https://ftp.ncbi.nlm.nih.gov/hapmap/phase_3/. To analyze the LP-associated allele and increase the density of variants in the Maasai, we imputed genotypes on Chromosome 2 in this population (from the HapMap Project Phase 3) using the TOPMed Imputation Server (version 3 reference panel) and retained imputed sites with r^2^ ≥ 0.9. The reference overlap with the Maasai was 97.68%.

