## Supplementary Data for "CLUES2 Companion: Computational pipelines to estimate, visualize, and date selection on multi-locus sites"

#### Supplementary Tables

**Table S1. Tabular output from CLUES2 Companion for the Bedouin population**. This Table shows the output of our Phase 2 pipeline in *.tsv format containing the rs identifiers for SNPs in the *MCM6* gene (including the rs41380347 SNP associated with lactase persistence), their genomic positions, the derived allele frequency at each SNP, logLR, -log10(p), epoch start, epoch end, *s* estimate (MLE1) and confidence interval.

**Table S2. Tabular output from CLUES2 Companion for the Maasai population**. Like Table S1, this Table shows the output of our Phase 2 pipeline in *.tsv format containing the rs identifiers for SNPs in the *MCM6* gene (including the rs145946881 SNP associated with lactase persistence), their genomic positions, the derived allele frequency at each SNP, logLR, -log10(p), epoch start, epoch end, *s* estimate (MLE1) and confidence interval.

**Table S3. Pairwise LD across the *MCM6* gene in the Finnish population**. We measured pairwise LD for SNPs across the ~36-kb region of *MCM6* using the $r^2$ statistic calculated with PLINK 2.0 (Chang et al. 2015) in the Finnish population The $r^2$ statistic, the square of the correlation coefficient, measure the non-random association between two indicator variables (in this case SNP loci). An $r^2=0$ means no association between loci while $r^2=1$ indicates perfect association; values between 0 and 1 signify degrees of association. In this Table, the chromosome number, rs number of the SNP associated with lactase persistence (rs4988235), the rs number of another SNP, their genomic positions, and $r^2$ are given.

**Table S4. Pairwise LD across the *MCM6* gene in the Bedouin population**. We measured pairwise LD for SNPs across the ~36-kb region of *MCM6* using the $r^2$ statistic calculated with PLINK 2.0 (Chang et al. 2015) in the Bedouin population. The $r^2$ statistic, the square of the correlation coefficient, measure the non-random association between two indicator variables (in this case SNP loci). An $r^2=0$ means no association between loci while $r^2=1$ indicates perfect association; values between 0 and 1 signify degrees of association. In this Table, the chromosome number, rs number of the SNP associated with lactase persistence (rs41380347), the rs number of another SNP, their genomic positions, and $r^2$ are given.

**Table S5. Pairwise LD across the *MCM6* gene in the Maasai population**. We measured pairwise LD for SNPs across the ~36-kb region of *MCM6* using the $r^2$ statistic calculated with PLINK 2.0 (Chang et al. 2015) in the Maasai population. The $r^2$ statistic, the square of the correlation coefficient, measures the non-random association between two indicator variables (in this case SNP loci). An $r^2=0$ means no association between loci while $r^2=1$ indicates perfect association; values between 0 and 1 signify degrees of association. In this Table, the chromosome number, rs number of the SNP associated with lactase persistence (rs41380347), the rs number of another SNP, their genomic positions, and $r^2$ are given.

**Table S6. Parameters of simulated and empirical data**. We simulated four datasets with known parameters for target SNPs (such as age of selection onset, *s* estimate, genomic position for target SNPs) using the sweep genic selection and sim_ancestry functions implemented in msprime (Baumdicker *et al.* 2022). Specifically, to generate the vcf files containing biallelic sites (0 represents the ancestral allele and 1 represents the derived), we applied the sweep genic selection function to create the genomic position of beneficial allele, the start and end frequencies of the beneficial allele, and the selection coefficient. Furthermore, we applied the sim_ancestry function to specify the number of samples, the model (sweep model), the population size, the recombination rate, and the length of the genomic region in these simulated datasets. **Panel A** shows a summary of the parameters used to simulate the data. We then applied our Phase 3 of our pipeline to these simulated data to see if our dating method would reproduce the ages of onset and the s estimates for the target SNPs with known genomic coordinates in the simulated data (**Panel B**). The overlap between key the simulated and empirical parameters of interest is shown in bold. The simulated data and associated scripts can be found on our repository at https://github.com/alisi1989/CLUES2-Companion/tree/main/example_sim.

#### Supplementary Figures

**Figure S1. Plot of s estimates for SNPs in the *MCM6* gene**. This plot shows estimates for SNPs (above a frequency of 5%) in the *MCM6* gene together with the rs identifiers for SNPs that exhibit significant s values *(P* < 0.05) for the Bedouin (**Panel A**) and the Maasai populations (**Panel B**) from the Middle East and East Africa, respectively. In the plot, each dot represents an *s* estimate, and the vertical bars indicate the confidence intervals around the estimate for each SNP. Furthermore, the color of each dot signifies the level of significance for each s estimate based on *P*–values in addition to the number of asterisks. The color scale for these dots is located at the extreme right-hand side of the plot.

Table S1. Tabular output from CLUES2 Companion for the Bedouin population

| rsID | POS | freq_der | logLR | -log10(p) | Epoch1_start | Epoch1_end | SelectionMLE1 | 95%_lower | 95%_upper |
|---|---|---|---|---|---|---|---|---|---|
| **rs55660827** | 135840873 | 0.2717 | 6.5549 | 3.53 | 0 | 536 | 0.09981 | 0.06556 | 0.13406 |
| **rs3820790** | 135841647 | 0.3370 | 1.9622 | 1.32 | 0 | 536 | 0.01848 | -0.00221 | 0.03917 |
| **rs55809728** | 135842606 | 0.3587 | 0.3093 | 0.36 | 0 | 536 | 0.00647 | -0.01226 | 0.02520 |
| **rs4988274** | 135843092 | 0.3152 | 4.8837 | 2.75 | 0 | 536 | 0.09981 | -0.02057 | 0.22018 |
| **rs4954490** | 135850661 | 0.2717 | 0.1389 | 0.22 | 0 | 536 | -0.01110 | -0.03956 | 0.01737 |
| **rs41380347** | 135851081 | 0.2717 | 6.5204 | 3.52 | 0 | 536 | 0.09981 | 0.06515 | 0.13448 |
| **rs4954493** | 135852405 | 0.2717 | 0.0055 | 0.04 | 0 | 536 | -0.00148 | -0.02431 | 0.02135 |
| **rs4988226** | 135853028 | 0.2500 | 3.6488 | 2.16 | 0 | 536 | 0.08443 | -0.00566 | 0.17451 |
| **rs309178** | 135854054 | 0.2500 | 3.1300 | 1.91 | 0 | 536 | 0.04741 | 0.00625 | 0.08858 |
| **rs309179** | 135856210 | 0.2500 | 3.9916 | 2.33 | 0 | 536 | 0.09987 | 0.01514 | 0.18461 |
| **rs309180** | 135856685 | 0.2500 | 3.0559 | 1.87 | 0 | 536 | 0.03465 | -0.00571 | 0.07500 |
| **rs309181** | 135857243 | 0.2500 | 3.4352 | 2.06 | 0 | 536 | 0.04495 | 0.00408 | 0.08582 |
| **rs61253125** | 135860608 | 0.3370 | 4.5626 | 2.60 | 0 | 536 | 0.04962 | 0.01245 | 0.08679 |
| **rs4988201** | 135860937 | 0.3370 | 4.0645 | 2.36 | 0 | 536 | 0.02131 | -0.00683 | 0.04944 |
| **rs309176** | 135864646 | 0.2500 | 2.8312 | 1.76 | 0 | 536 | 0.02584 | -0.00904 | 0.06071 |
| **rs3087343** | 135864973 | 0.3370 | 4.2954 | 2.47 | 0 | 536 | 0.02593 | -0.00036 | 0.05221 |
| **rs4594504** | 135867543 | 0.3370 | 4.2370 | 2.44 | 0 | 536 | 0.02931 | 0.00389 | 0.05473 |
| **rs309811** | 135868032 | 0.2500 | 3.2024 | 1.94 | 0 | 536 | 0.04309 | 0.00908 | 0.07709 |
| **rs3769001** | 135868508 | 0.3370 | 4.6018 | 2.62 | 0 | 536 | 0.04659 | 0.01490 | 0.07827 |
| **rs4988163** | 135870551 | 0.3370 | 4.1360 | 2.40 | 0 | 536 | 0.03279 | 0.00267 | 0.06290 |
| **rs309128** | 135872341 | 0.2500 | 3.3569 | 2.02 | 0 | 536 | 0.08035 | -0.00625 | 0.16695 |
| **rs680428** | 135873187 | 0.2500 | 2.8326 | 1.76 | 0 | 536 | 0.02701 | -0.00681 | 0.06083 |
| **rs188680** | 135873419 | 0.2500 | 2.8300 | 1.76 | 0 | 536 | 0.03143 | 0.00000 | 0.06285 |
| **rs309130** | 135873461 | 0.2500 | 3.0688 | 1.88 | 0 | 536 | 0.03898 | 0.00813 | 0.06983 |
| **rs4988145** | 135874730 | 0.3261 | 4.0015 | 2.33 | 0 | 536 | 0.03211 | 0.00269 | 0.06154 |

Table S2. Tabular output from CLUES2 Companion for the Maasai population

| rsID | POS | der_freq | logLR | -log10(p) | Epoch1_start | Epoch1_end | SelectionMLE1 | 95%_lower | 95%_upper |
|---|---|---|---|---|---|---|---|---|---|
| **rs4988285** | 135839996 | 0.0625 | 2.0629 | 1.37 | 0 | 536 | -0.01589 | -0.02695 | -0.00484 |
| **rs3769006** | 135841440 | 0.0625 | 2.3440 | 1.52 | 0 | 536 | -0.01732 | -0.02824 | -0.00641 |
| **rs4988269** | 135844721 | 0.0598 | 1.4854 | 1.07 | 0 | 536 | 0.02887 | -0.02851 | 0.08626 |
| **rs3754686** | 135845706 | 0.0571 | 1.3680 | 1.01 | 0 | 536 | 0.02198 | -0.03444 | 0.07841 |
| **rs3769004** | 135846020 | 0.0625 | 2.1424 | 1.42 | 0 | 536 | -0.01629 | -0.02747 | -0.00511 |
| **rs4988265** | 135846068 | 0.0897 | 3.4382 | 2.06 | 0 | 536 | -0.01681 | -0.02590 | -0.00773 |
| **rs3769003** | 135846120 | 0.0897 | 2.7954 | 1.74 | 0 | 536 | -0.01539 | -0.02477 | -0.00601 |
| **rs4988262** | 135846640 | 0.0815 | 2.6984 | 1.70 | 0 | 536 | -0.01512 | -0.02478 | -0.00545 |
| **rs4988240** | 135850362 | 0.0625 | 1.3375 | 0.99 | 0 | 536 | 0.02947 | -0.04497 | 0.10390 |
| **rs4954490** | 135850661 | 0.0625 | 1.4875 | 1.07 | 0 | 536 | 0.02159 | -0.03402 | 0.07720 |
| **rs145946881** | 135851176 | 0.5462 | 18.1547 | 8.77 | 0 | 536 | 0.09981 | 0.08799 | 0.11163 |
| **rs4954493** | 135852405 | 0.0652 | 1.3837 | 1.02 | 0 | 536 | 0.02054 | -0.05308 | 0.09416 |
| **rs4988226** | 135853028 | 0.0571 | 1.3680 | 1.01 | 0 | 536 | 0.02198 | -0.03444 | 0.07841 |
| **rs309178** | 135854054 | 0.0571 | 1.2555 | 0.95 | 0 | 536 | 0.02333 | -0.02674 | 0.07341 |
| **rs111837148** | 135854466 | 0.6957 | 15.4124 | 7.55 | 0 | 536 | 0.06048 | 0.03643 | 0.08454 |
| **rs112728941** | 135855028 | 0.0897 | 3.2612 | 1.97 | 0 | 536 | -0.01644 | -0.02571 | -0.00717 |
| **rs309179** | 135856210 | 0.0625 | 1.3770 | 1.01 | 0 | 536 | 0.02610 | -0.03492 | 0.08712 |
| **rs309180** | 135856685 | 0.0571 | 1.1909 | 0.91 | 0 | 536 | 0.01926 | -0.04902 | 0.08753 |
| **rs309181** | 135857243 | 0.0652 | 1.5734 | 1.12 | 0 | 536 | 0.03184 | -0.02612 | 0.08980 |
| **rs4988205** | 135858466 | 0.0625 | 2.4695 | 1.58 | 0 | 536 | -0.01759 | -0.02837 | -0.00681 |
| **rs309173** | 135859954 | 0.0625 | 1.4187 | 1.04 | 0 | 536 | 0.02239 | -0.03179 | 0.07656 |
| **rs111427863** | 135860347 | 0.0598 | 1.3760 | 1.01 | 0 | 536 | 0.03881 | -0.02901 | 0.10663 |
| **rs61253125** | 135860608 | 0.6957 | 18.1697 | 8.78 | 0 | 536 | 0.09984 | 0.07429 | 0.12539 |
| **rs4988201** | 135860937 | 0.6957 | 19.7381 | 9.48 | 0 | 536 | 0.09984 | 0.09145 | 0.10824 |
| **rs75597312** | 135862045 | 0.0598 | 1.3238 | 0.98 | 0 | 536 | 0.02775 | -0.04270 | 0.09819 |
| **rs4988184** | 135864398 | 0.0897 | 3.4382 | 2.06 | 0 | 536 | -0.01681 | -0.02590 | -0.00773 |
| **rs309176** | 135864646 | 0.0571 | 1.4741 | 1.07 | 0 | 536 | 0.01880 | -0.03679 | 0.07438 |
| **rs3087343** | 135864973 | 0.6957 | 17.1677 | 8.33 | 0 | 536 | 0.09935 | 0.05530 | 0.14341 |
| **rs309177** | 135865255 | 0.0788 | 0.7152 | 0.64 | 0 | 536 | 0.01448 | -0.00427 | 0.03323 |
| **rs2289048** | 135866330 | 0.0897 | 2.8252 | 1.76 | 0 | 536 | -0.01594 | -0.02563 | -0.00626 |
| **rs2289049** | 135866553 | 0.0897 | 2.7726 | 1.73 | 0 | 536 | -0.01552 | -0.02509 | -0.00596 |
| **rs2070068** | 135866585 | 0.0897 | 3.3192 | 2.00 | 0 | 536 | -0.01697 | -0.02626 | -0.00767 |
| **rs1435577** | 135867377 | 0.7011 | 20.0135 | 9.60 | 0 | 536 | 0.09986 | 0.09160 | 0.10811 |
| **rs4594504** | 135867543 | 0.6957 | 14.7583 | 7.26 | 0 | 536 | 0.05099 | 0.02159 | 0.08038 |
| **rs3769001** | 135868508 | 0.6929 | 17.2316 | 8.36 | 0 | 536 | 0.07536 | 0.05092 | 0.09980 |
| **rs4988166** | 135869937 | 0.0598 | 1.4034 | 1.03 | 0 | 536 | 0.03012 | -0.03779 | 0.09804 |
| **rs4988163** | 135870551 | 0.6929 | 15.9788 | 7.80 | 0 | 536 | 0.08754 | 0.05013 | 0.12495 |
| **rs4988161** | 135871101 | 0.0897 | 2.6584 | 1.68 | 0 | 536 | -0.01531 | -0.02505 | -0.00556 |
| **rs309128** | 135872341 | 0.0598 | 1.3997 | 1.03 | 0 | 536 | 0.02220 | -0.03075 | 0.07515 |
| **rs4988158** | 135872417 | 0.0598 | 1.3040 | 0.97 | 0 | 536 | 0.03335 | -0.04124 | 0.10795 |

### Table S2

| | | | | | | | | | |
|---|---|---|---|---|---|---|---|---|---|
| **rs680428** | 135873187 | 0.0598 | 1.2940 | 0.97 | 0 | 536 | 0.02371 | -0.02837 | 0.07579 |
| **rs188680** | 135873419 | 0.0598 | 1.2500 | 0.94 | 0 | 536 | 0.02241 | -0.03911 | 0.08394 |
| **rs309130** | 135873461 | 0.0571 | 1.4631 | 1.06 | 0 | 536 | 0.01898 | -0.02325 | 0.06121 |
| **rs4988145** | 135874730 | 0.6957 | 16.1052 | 7.86 | 0 | 536 | 0.08775 | 0.05086 | 0.12463 |
| **rs309132** | 135875441 | 0.7663 | 1.5779 | 1.12 | 0 | 536 | 0.01035 | 0.00057 | 0.02013 |

Table S3. Pairwise LD across the *MCM6* gene in the Finnish population

| CHR | POS_SNP1 | rs_SNP1 | POS_SNP2 | rs_SNP2 | r2 |
|---|---|---|---|---|---|
| 2 | 135841647 | rs3820790 | 135851076 | rs4988235 | 0.254395 |
| 2 | 135842606 | rs55809728 | 135851076 | rs4988235 | 0.0740056 |
| 2 | 135843092 | rs4988274 | 135851076 | rs4988235 | 0.200302 |
| 2 | 135844921 | rs3739020 | 135851076 | rs4988235 | 0.698447 |
| 2 | 135845706 | rs3754686 | 135851076 | rs4988235 | 0.756734 |
| 2 | 135845796 | rs3769005 | 135851076 | rs4988235 | 0.756734 |
| 2 | 135850133 | rs4988243 | 135851076 | rs4988235 | 0.173947 |
| 2 | 135850661 | rs4954490 | 135851076 | rs4988235 | 0.756734 |
| 2 | 135851076 | rs4988235 | 135852405 | rs4954493 | 0.756734 |
| 2 | 135851076 | rs4988235 | 135853028 | rs4988226 | 0.756734 |
| 2 | 135851076 | rs4988235 | 135854054 | rs309178 | 0.756734 |
| 2 | 135851076 | rs4988235 | 135854466 | rs111837148 | 0.196312 |
| 2 | 135851076 | rs4988235 | 135856210 | rs309179 | 0.756734 |
| 2 | 135851076 | rs4988235 | 135856685 | rs309180 | 0.756734 |
| 2 | 135851076 | rs4988235 | 135857243 | rs309181 | 0.756734 |
| 2 | 135851076 | rs4988235 | 135857652 | rs3213871 | 0.306552 |
| 2 | 135851076 | rs4988235 | 135859184 | rs182549 | 1 |
| 2 | 135851076 | rs4988235 | 135859907 | rs12474093 | 0.191563 |
| 2 | 135851076 | rs4988235 | 135859954 | rs309173 | 0.756734 |
| 2 | 135851076 | rs4988235 | 135860235 | rs160329 | 0.87168 |
| 2 | 135851076 | rs4988235 | 135860608 | rs61253125 | 0.196312 |
| 2 | 135851076 | rs4988235 | 135860937 | rs4988201 | 0.196312 |
| 2 | 135851076 | rs4988235 | 135864476 | rs4988183 | 0.698447 |
| 2 | 135851076 | rs4988235 | 135864646 | rs309176 | 0.756734 |
| 2 | 135851076 | rs4988235 | 135864973 | rs3087343 | 0.196312 |
| 2 | 135851076 | rs4988235 | 135867349 | rs56263017 | 0.306552 |
| 2 | 135851076 | rs4988235 | 135867377 | rs1435577 | 0.196312 |
| 2 | 135851076 | rs4988235 | 135867543 | rs4594504 | 0.196312 |
| 2 | 135851076 | rs4988235 | 135868032 | rs309811 | 0.756734 |

| | | | | | |
|---|---|---|---|---|---|
| 2 | 135851076 | rs4988235 | 135868434 | rs3769002 | 0.306552 |
| 2 | 135851076 | rs4988235 | 135868508 | rs3769001 | 0.196312 |
| 2 | 135851076 | rs4988235 | 135870551 | rs4988163 | 0.196312 |
| 2 | 135851076 | rs4988235 | 135870810 | rs4954513 | 0.306552 |
| 2 | 135851076 | rs4988235 | 135872341 | rs309128 | 0.756734 |
| 2 | 135851076 | rs4988235 | 135873187 | rs680428 | 0.756734 |
| 2 | 135851076 | rs4988235 | 135873419 | rs188680 | 0.756734 |
| 2 | 135851076 | rs4988235 | 135873461 | rs309130 | 0.756734 |
| 2 | 135851076 | rs4988235 | 135874730 | rs4988145 | 0.196312 |
| 2 | 135851076 | rs4988235 | 135875441 | rs309132 | 0.306552 |
| 2 | 135851076 | rs4988235 | 135876201 | rs191079 | 0.756734 |
| 2 | 135851076 | rs4988235 | 135876392 | rs1057031 | 0.198227 |

Table S4. Pairwise LD across the *MCM6* gene in the Bedouin population

| CHR | POS_SNP1 | rs_SNP1 | POS_SNP2 | rs_SNP2 | r2 |
|---|---|---|---|---|---|
| 2 | 135840873 | rs55660827 | 135851081 | rs41380347 | 1 |
| 2 | 135841647 | rs3820790 | 135851081 | rs41380347 | 0.800944 |
| 2 | 135842606 | rs55809728 | 135851081 | rs41380347 | 0.748572 |
| 2 | 135843092 | rs4988274 | 135851081 | rs41380347 | 0.151676 |
| 2 | 135845706 | rs3754686 | 135851081 | rs41380347 | 0.149461 |
| 2 | 135845796 | rs3769005 | 135851081 | rs41380347 | 0.16993 |
| 2 | 135850133 | rs4988243 | 135851081 | rs41380347 | 0.829507 |
| 2 | 135850661 | rs4954490 | 135851081 | rs41380347 | 0.149461 |
| 2 | 135851081 | rs41380347 | 135852405 | rs4954493 | 0.149461 |
| 2 | 135851081 | rs41380347 | 135853028 | rs4988226 | 0.113509 |
| 2 | 135851081 | rs41380347 | 135854054 | rs309178 | 0.113509 |
| 2 | 135851081 | rs41380347 | 135856210 | rs309179 | 0.113509 |
| 2 | 135851081 | rs41380347 | 135856685 | rs309180 | 0.113509 |
| 2 | 135851081 | rs41380347 | 135857243 | rs309181 | 0.113509 |
| 2 | 135851081 | rs41380347 | 135857652 | rs3213871 | 0.702103 |
| 2 | 135851081 | rs41380347 | 135860608 | rs61253125 | 0.192068 |
| 2 | 135851081 | rs41380347 | 135860937 | rs4988201 | 0.192068 |
| 2 | 135851081 | rs41380347 | 135864646 | rs309176 | 0.113509 |
| 2 | 135851081 | rs41380347 | 135864973 | rs3087343 | 0.192068 |
| 2 | 135851081 | rs41380347 | 135867349 | rs56263017 | 0.702103 |
| 2 | 135851081 | rs41380347 | 135867543 | rs4594504 | 0.192068 |
| 2 | 135851081 | rs41380347 | 135868032 | rs309811 | 0.113509 |
| 2 | 135851081 | rs41380347 | 135868434 | rs3769002 | 0.702103 |
| 2 | 135851081 | rs41380347 | 135868508 | rs3769001 | 0.192068 |
| 2 | 135851081 | rs41380347 | 135870551 | rs4988163 | 0.192068 |
| 2 | 135851081 | rs41380347 | 135872341 | rs309128 | 0.113509 |
| 2 | 135851081 | rs41380347 | 135873187 | rs680428 | 0.113509 |
| 2 | 135851081 | rs41380347 | 135873419 | rs188680 | 0.113509 |
| 2 | 135851081 | rs41380347 | 135873461 | rs309130 | 0.113509 |

| | | | | | |
|---|---|---|---|---|---|
| 2 | 135851081 | rs41380347 | 135874730 | rs4988145 | 0.186236 |
| 2 | 135851081 | rs41380347 | 135875441 | rs309132 | 0.604293 |
| 2 | 135851081 | rs41380347 | 135876201 | rs191079 | 0.113509 |
| 2 | 135851081 | rs41380347 | 135876392 | rs1057031 | 0.192068 |

Table S5. Pairwise LD across the *MCM6* gene in the Maasai population

| CHR | POS_SNP1 | rs_SNP1 | POS_SNP2 | rs_SNP2 | r2 |
|---|---|---|---|---|---|
| 2 | 135839996 | rs4988285 | 135851176 | rs145946881 | 0.0855107 |
| 2 | 135841440 | rs3769006 | 135851176 | rs145946881 | 0.0855107 |
| 2 | 135844721 | rs4988269 | 135851176 | rs145946881 | 0.0874969 |
| 2 | 135844921 | rs3739020 | 135851176 | rs145946881 | 0.351705 |
| 2 | 135845706 | rs3754686 | 135851176 | rs145946881 | 0.0561369 |
| 2 | 135845796 | rs3769005 | 135851176 | rs145946881 | 0.0911562 |
| 2 | 135846020 | rs3769004 | 135851176 | rs145946881 | 0.0855107 |
| 2 | 135846068 | rs4988265 | 135851176 | rs145946881 | 0.0978881 |
| 2 | 135846120 | rs3769003 | 135851176 | rs145946881 | 0.0978881 |
| 2 | 135846640 | rs4988262 | 135851176 | rs145946881 | 0.0884191 |
| 2 | 135846775 | rs4988261 | 135851176 | rs145946881 | 0.0978881 |
| 2 | 135846776 | rs4988260 | 135851176 | rs145946881 | 0.0978881 |
| 2 | 135847541 | rs74266308 | 135851176 | rs145946881 | 0.0978881 |
| 2 | 135850362 | rs4988240 | 135851176 | rs145946881 | 0.116047 |
| 2 | 135850661 | rs4954490 | 135851176 | rs145946881 | 0.0777809 |
| 2 | 135851176 | rs145946881 | 135852405 | rs4954493 | 0.0894183 |
| 2 | 135851176 | rs145946881 | 135853028 | rs4988226 | 0.0561369 |
| 2 | 135851176 | rs145946881 | 135854054 | rs309178 | 0.0561369 |
| 2 | 135851176 | rs145946881 | 135854466 | rs111837148 | 0.507701 |
| 2 | 135851176 | rs145946881 | 135855028 | rs112728941 | 0.0978881 |
| 2 | 135851176 | rs145946881 | 135856210 | rs309179 | 0.0777809 |
| 2 | 135851176 | rs145946881 | 135856685 | rs309180 | 0.0561369 |
| 2 | 135851176 | rs145946881 | 135857243 | rs309181 | 0.0894183 |
| 2 | 135851176 | rs145946881 | 135857385 | rs4988211 | 0.0978881 |
| 2 | 135851176 | rs145946881 | 135857652 | rs3213871 | 0.113376 |
| 2 | 135851176 | rs145946881 | 135858466 | rs4988205 | 0.0855107 |
| 2 | 135851176 | rs145946881 | 135859954 | rs309173 | 0.0777809 |
| 2 | 135851176 | rs145946881 | 135860347 | rs111427863 | 0.0874969 |
| 2 | 135851176 | rs145946881 | 135860608 | rs61253125 | 0.507701 |

| | | | | | |
|---|---|---|---|---|---|
| 2 | 135851176 | rs145946881 | 135860937 | rs4988201 | 0.507701 |
| 2 | 135851176 | rs145946881 | 135861544 | rs4988196 | 0.0978881 |
| 2 | 135851176 | rs145946881 | 135861643 | rs4988194 | 0.0978881 |
| 2 | 135851176 | rs145946881 | 135862045 | rs75597312 | 0.0874969 |
| 2 | 135851176 | rs145946881 | 135864398 | rs4988184 | 0.0978881 |
| 2 | 135851176 | rs145946881 | 135864476 | rs4988183 | 0.351705 |
| 2 | 135851176 | rs145946881 | 135864646 | rs309176 | 0.0561369 |
| 2 | 135851176 | rs145946881 | 135864973 | rs3087343 | 0.507701 |
| 2 | 135851176 | rs145946881 | 135865255 | rs309177 | 0.0896627 |
| 2 | 135851176 | rs145946881 | 135866330 | rs2289048 | 0.0978881 |
| 2 | 135851176 | rs145946881 | 135866553 | rs2289049 | 0.0978881 |
| 2 | 135851176 | rs145946881 | 135866585 | rs2070068 | 0.0978881 |
| 2 | 135851176 | rs145946881 | 135867349 | rs56263017 | 0.113376 |
| 2 | 135851176 | rs145946881 | 135867377 | rs1435577 | 0.497349 |
| 2 | 135851176 | rs145946881 | 135867543 | rs4594504 | 0.507701 |
| 2 | 135851176 | rs145946881 | 135868434 | rs3769002 | 0.113376 |
| 2 | 135851176 | rs145946881 | 135868508 | rs3769001 | 0.528781 |
| 2 | 135851176 | rs145946881 | 135869937 | rs4988166 | 0.0874969 |
| 2 | 135851176 | rs145946881 | 135870551 | rs4988163 | 0.528781 |
| 2 | 135851176 | rs145946881 | 135870810 | rs4954513 | 0.113376 |
| 2 | 135851176 | rs145946881 | 135871101 | rs4988161 | 0.0978881 |
| 2 | 135851176 | rs145946881 | 135872341 | rs309128 | 0.0666736 |
| 2 | 135851176 | rs145946881 | 135872417 | rs4988158 | 0.0874969 |
| 2 | 135851176 | rs145946881 | 135872600 | rs3739019 | 0.0978881 |
| 2 | 135851176 | rs145946881 | 135873187 | rs680428 | 0.0666736 |
| 2 | 135851176 | rs145946881 | 135873419 | rs188680 | 0.0666736 |
| 2 | 135851176 | rs145946881 | 135873461 | rs309130 | 0.0561369 |
| 2 | 135851176 | rs145946881 | 135873845 | rs3769000 | 0.124749 |
| 2 | 135851176 | rs145946881 | 135874555 | rs73957046 | 0.121352 |
| 2 | 135851176 | rs145946881 | 135874730 | rs4988145 | 0.507701 |

| | | | | | |
|---|---|---|---|---|---|
| 2 | 135851176 | rs145946881 | 135875441 | rs309132 | 0.360993 |
| 2 | 135851176 | rs145946881 | 135876392 | rs1057031 | 0.479267 |

Table S6. Parameters of simulated and empirical data

**A**

**Theoretical Simulation Parameters**

| | Simulation 1 | Simulation 2 | Simulation 3 | Simulation 4 |
|---|---|---|---|---|
| Effective population size (Ne) | 20,000 | 20,000 | 20,000 | 20,000 |
| Sequence length | 10,000,000 | 10,000,000 | 10,000,000 | 10,000,000 |
| Recombination rate | 0.00000001 | 0.00000001 | 0.00000001 | 0.00000001 |
| Mutation rate | 0.0000000125 | 0.0000000125 | 0.0000000125 | 0.0000000125 |
| s estimate | **0.02187** | **0.02187** | **0.02187** | **0.02187** |
| Genomic position | 1,500,000 | 1,500,000 | 1,500,000 | 1,500,000 |
| Number of individuals | 100 | 100 | 100 | 100 |
| Theoretical Onset of selection | **129.6** | **240.0** | **330.1** | **429.4** |

**B**

**Empirical Parameters**

| | CLUES2 Companion 1 | CLUES2 Companion 2 | CLUES2 Companion 3 | CLUES2 Companion 4 |
|---|---|---|---|---|
| Effective population size (Ne) | 20,000 | 20,000 | 20,000 | 20,000 |
| Sequence length | 10,000,000 | 10,000,000 | 10,000,000 | 10,000,000 |
| Recombination rate | 0.00000001 | 0.00000001 | 0.00000001 | 0.00000001 |
| Mutation rate | 0.0000000125 | 0.0000000125 | 0.0000000125 | 0.0000000125 |
| s estimate | **0.02187** | **0.02187** | **0.02187** | **0.02187** |
| Genomic position | 1,500,000 | 1,500,000 | 1,500,000 | 1,500,000 |
| Number of individuals | 100 | 100 | 100 | 100 |
| Theoretical Onset of selection | **125** | **225** | **325** | **425** |

A

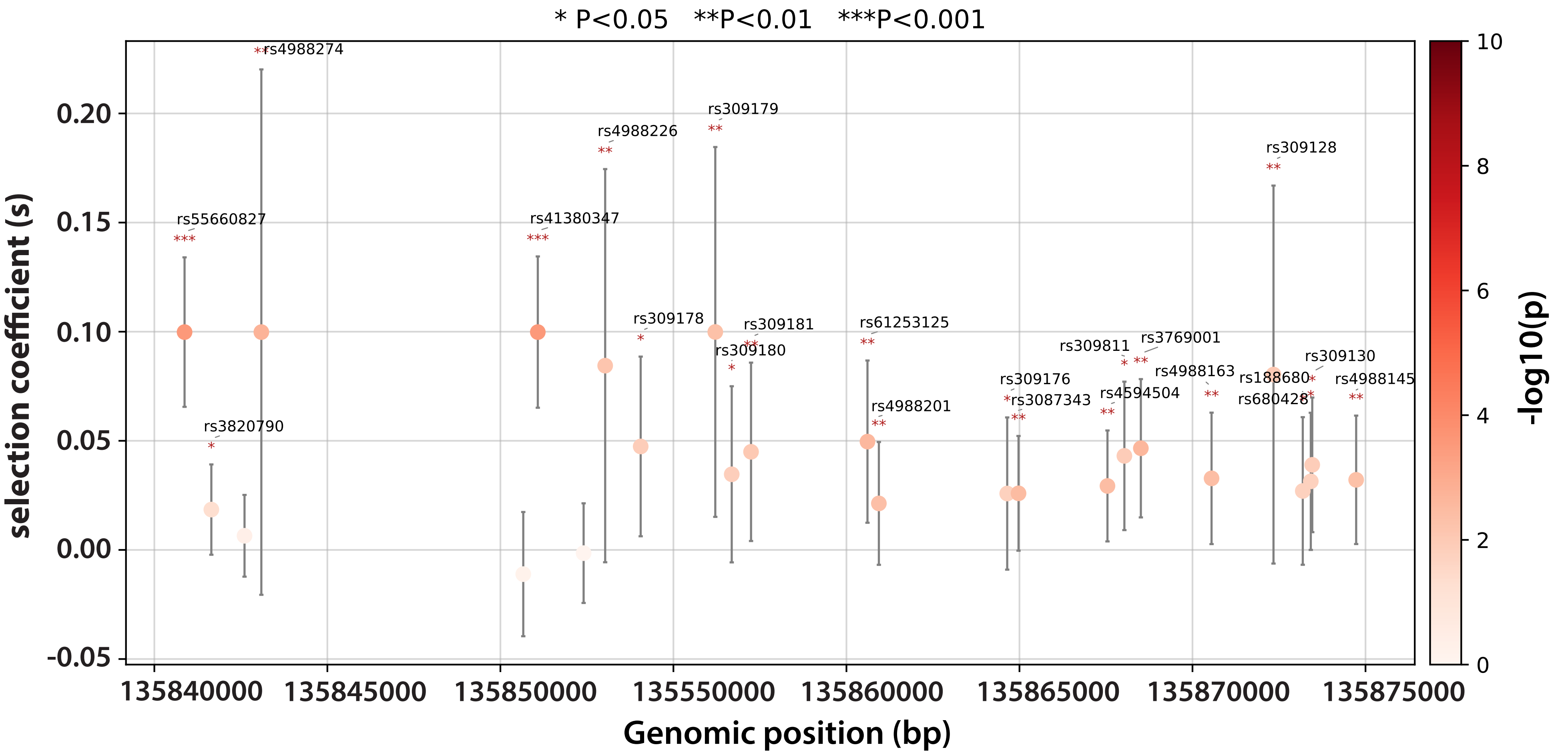

* P<0.05 **P<0.01 ***P<0.001
selection coefficient (s)
Genomic position (bp)
-log10(p)
rs4988274
rs55660827
rs3820790
rs41380347
rs4988226
rs309179
rs309178
rs309181
rs309180
rs61253125
rs4988201
rs309176
rs3087343
rs309811
rs3769001
rs4594504
rs4988163
rs309128
rs188680
rs680428
rs309130
rs4988145


B

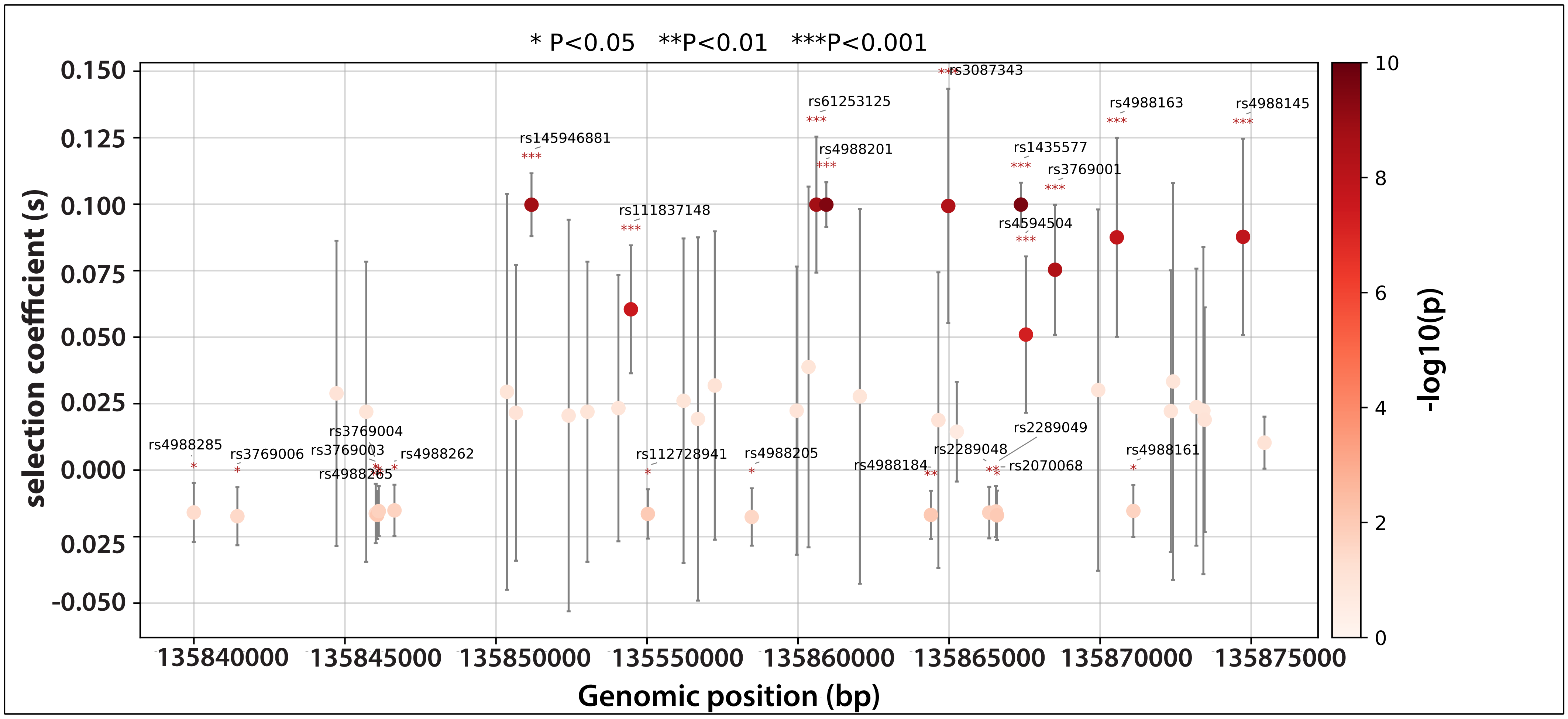

* P<0.05 **P<0.01 ***P<0.001
selection coefficient (s)
Genomic position (bp)
-log10(p)
rs145946881
rs111837148
rs61253125
rs4988201
rs3087343
rs1435577
rs3769001
rs4594504
rs4988163
rs4988145
rs4988285
rs3769006
rs3769004
rs3769003
rs4988265
rs4988262
rs112728941
rs4988205
rs4988184
rs2289048
rs2289049
rs2070068
rs4988161


**Figure S1**
